# Cardiac microtubules mediate transverse (t)-tubule growth and homeostasis

**DOI:** 10.64898/2026.08.16.745070

**Authors:** A.S. Whitley, G.W.P. Madders, A. Livesey, A. Ashik, K. Uchida, B. Prosser, A.W.T. Trafford, K.M. Dibb

## Abstract

Transverse (t)-tubules enable rapid, synchronous Ca release required for efficient cardiac contraction by bringing L-type Ca channels into close apposition with ryanodine receptors. In heart failure with reduced ejection fraction (HFrEF), t-tubule disorganisation and loss occur alongside cardiac microtubule remodelling, contributing to impaired Ca handling and contractile dysfunction. Despite their canonical function in contraction, how t-tubules develop is unknown. Microtubules support delivery of L-type Ca channels to t-tubules via Amphiphysin-II/BIN1, yet whether microtubules directly regulate t-tubule formation and maintenance is unclear. Here, we investigated a role for microtubules in t-tubule development and homeostasis.

Neonatal rat ventricular myocytes (NRVMs), which lack endogenous t-tubules, were used as a reductionist model in which BIN1 overexpression induces nascent membrane tubules. Microtubule depolymerisation with nocodazole before BIN1 overexpression impaired BIN1-driven tubule formation, reducing tubule density and length. Dynein inhibition with EHNA produced similar effects, indicating a requirement for microtubule-based motor activity during tubule elongation. Knockdown of the microtubule +TIP tracking protein CLIP-170 also reduced BIN1-driven tubule density, implicating BIN1–CLIP-170-dependent microtubule capture in tubule initiation.

Microtubules were also required to maintain existing tubules. In NRVMs with established BIN1-driven tubules, microtubule depolymerisation, microtubule stabilisation or dynein inhibition each reduced tubule density and length. Consistent with this, acute microtubule depolymerisation or stabilisation disrupted native t-tubule networks in isolated adult sheep left atrial myocytes.

Together, these findings identify cardiac microtubules as active regulators of t-tubule architecture. We propose that BIN1-dependent tubule formation requires CLIP-170-mediated microtubule plus-end capture and dynein-dependent elongation, while ongoing microtubule dynamics are necessary to preserve mature t-tubule structure.

## Introduction

Cardiac transverse (t)-tubules are deep, perpendicular invaginations of the cardiac myocyte surface membrane that underlie the rapid and synchronous rise in Ca^2+^ necessary for healthy, forceful contraction [1–4]. T-tubules enable the functional coupling of the L-type Ca^2+^ channel (LTCC), situated primarily on t-tubule membranes, and ryanodine receptors (cardiac isoform RyR2) residing upon the sarcoplasmic reticulum (SR) membrane [5–8]. The close proximity (∼12 nm) of the channels that form the dyad allows rapid Ca^2+^ entry via LTCCs upon membrane depolarisation to trigger RyR Ca^2+^ release from the SR in a process termed Ca^2+^-induced Ca^2+^-release (CICR) [9–12]. Myocyte architecture therefore underpins rapid, synchronous contraction and systolic function. The formation and maintenance of this specialised membrane architecture both depend not only on structural support but also on intracellular trafficking processes mediated by the microtubule cytoskeleton.

Cardiac microtubules are long, hollow and highly dynamic polymers of α/β-tubulin that extend throughout the cardiomyocyte cytoplasm and form a major component of the non-sarcomeric cytoskeleton [13, 14]. In addition to acting as intracellular ‘train tracks’, microtubules contribute directly to cardiac mechanics by buckling during systole and bearing compressive load [15, 16], and provide a platform for compartmentalised signalling and cargo delivery by mediating stretch-dependent X-ROS signalling [17], mRNA/ribosome localisation and local translation during cardiac growth [18, 19], and spatially restricted β-adrenergic signalling domains [20, 21]. Importantly for t-tubule biology, BIN1-rich membrane scaffolds can tether dynamic microtubules and direct microtubule-dependent delivery of LTCCs to t-tubule membranes [22–24], providing a mechanistic link between the microtubule cytoskeleton, dyadic organisation and t-tubules.

We and others have previously shown that in heart failure with reduced ejection fraction (HFrEF), ventricular t-tubules are decreased in density and atrial t-tubules are almost entirely lost, together with a reduction in *I_CaL_*, orphaned RyRs and contractile dysfunction, resulting in a reduction in Ca^2+^ transient amplitude and perpetuating systolic dysfunction [2, 25–33]. Concomitantly, ventricular microtubules densify and impede load during systole, perpetuating contractile dysfunction further in HFrEF [34–37]. However, t-tubules and microtubules are labile structures which rapidly turnover. Given our recent work in tachypacing-induced heart failure has shown that functional t-tubules can be recovered when tachypacing is ceased alongside Ca^2+^ homeostasis [38], we hypothesised that microtubules may play a role in t-tubule development and turnover such that disruption in one system may affect the other.

Initial cardiac t-tubule invaginations are thought to form from BIN1-rich membrane scaffolds that are subsequently driven inwards via a dual mechanism involving the accrual of membrane lipids and t-tubule forming proteins, alongside the fusion of pre-formed vesicle tandems containing LTCCs and RyRs into early cardiac dyads [39–41]. This process is also regulated by key structural proteins, myotubularin (MTM1) and junctophilin-2 (JPH2), which promote t-tubule membrane curvature, stabilisation and maturation of dyadic Ca²⁺ release units [23, 42–45]. However, in the atria specifically, postnatal t-tubule development is species-dependent, the mature network being sparse or largely absent in the atria of small mammals but well developed in large mammals, including sheep and humans [2, 46–49]. As atrial t-tubules develop, they establish internal dyadic Ca²⁺ release sites by bringing LTCCs into close apposition with RyR2 clusters, thereby converting immature, peripheral Ca²⁺ propagation into rapid, spatially synchronised Ca²⁺ transients that support coordinated contraction [48, 50].

Post-formation, t-tubule homeostasis is also a continuous process, as t-tubules have been shown to rapidly turnover to allow adaptation to stressors such as exercise, mechanical unloading, transaortic constriction (TAC) [32, 51–54]. In non-cardiac cells, microtubules have been shown to influence the formation and turnover of endocytic tubules via the molecular microtubule motor protein dynein, which walks along microtubules in a retrograde (anterior) motion and ‘pulls’ membrane-associated endocytic tubules inwards [55]. In HeLa cells, microtubule depolymerisation with nocodazole decreased BIN1-driven tubule density via reduced BIN1: CLIP-170 interactions at the microtubule +TIP, and in cardiac myocytes, microtubules are shown to dynamically align with t-tubules during homeostasis [56, 57]. Given that BIN1-dependent membrane remodelling and LTCC delivery are both linked to the microtubule cytoskeleton, microtubules are well positioned to coordinate the trafficking and forces required for t-tubule development. However, in both atria and ventricles, whether cardiac microtubules drive t-tubule biogenesis or play an important role in t-tubule homeostasis remains to be determined.

Here, we test the hypothesis that cardiac microtubules provide the structural tracks and motors required for BIN1-driven tubule formation and ongoing t-tubule homeostasis. Using BIN1-induced tubule formation in neonatal rat ventricular myocytes and native t-tubules in adult sheep atrial myocytes, we define a microtubule-dependent mechanism involving CLIP-170 anchoring, dynein-driven elongation and dynamic microtubules. These findings identify cardiac microtubules as active regulators of t-tubule architecture, rather than passive cytoskeletal bystanders.

## Methods

### Ethical Approval

All experiments were conducted according to the UK Animals (Scientific Procedures) Act, 1986, and local ethical approval was obtained from The University of Manchester Animal Welfare and Ethical Review Board. Reporting of animal experiments was in accordance with the ARRIVE guidelines 2.0.

### Husbandry

Female Welsh Mountain sheep were provided by the Biological Services Facility, The University of Manchester. Power calculations were performed, after determination of between and within subject variance, based on preliminary data or previous publications. Group sizes are given in figure legends and were determined using α = 0.05 and 1-b = 0.8. Animals were housed on a 12-hour light: dark cycle and fed hay and water *ad libitum*.

### Isolation of atrial myocytes

Single left atrial myocytes were isolated using a langendorff collagenase and protease digestion technique as described previously [2]. In brief, animals were euthanised with an intravenous overdose of pentobarbitone (200 mg/kg). Heparin (10,000 U) was administered prior to prevent coagulation. The heart was rapidly excised and the left atrial circumflex artery was cannulated to perfuse the left atria. Individual left atrial myocytes were isolated using collagenase (Worthington Type II, 0.1 mg/ml) and protease (from *Streptomyces griseus*, 0.02 mg/ml) dissolved in an isolation solution containing (in mmol/l; NaCl, 134; glucose, 11; HEPES, 10; BDM, 10; KCl, 4; MgSO_4_, 1.2; Na_2_HPO_4_, 1.2; and BSA, 0.5 mg.ml^−1^, pH 7.34 with NaOH). Once visual signs of digestion occurred, the enzyme was removed and the atria was perfused for 20 minutes in a taurine solution containing (in mmol/l; NaCl, 113; taurine, 50; glucose, 11; HEPES, 10; BDM, 10; KCl, 4; MgSO_4_, 1.2; Na_2_HPO_4_, 1.2; CaCl_2_, 0.1 and BSA, 0.5 mg.ml^−1^, pH 7.34 with NaOH) before being roughly dissected and then dispersed by gentle trituration.

### Isolation of Neonatal Rat Ventricular Myocytes

Neonatal rat ventricular myocytes (NRVMs) were isolated from entire litters of 2–3-day old Wistar rat pups and plated onto 8-well plates (Ibidi µ-plates; Thistle Scientific UK) at a density of ∼3 x 10^5^ cells/ml in plating medium (DMEM, 68%; M199, 17%; normal horse serum, 10%; fetal bovine serum, 5%). In brief, pups were culled by cervical dislocation, atria and fatty tissues removed, before ventricles were rapidly excised and placed in ice-cold dissociation buffer (mmol/l: NaCl, 116; HEPES, 20; glucose, 5.6; KCl, 5.4; NaH_2_PO_4,_ 1; MgSO_4_, 0.83; pH 7.35). Single NRVMs were enzymatically digested by gentle stirring (120 rpm, 7 minutes, 37°C) and trituration, using 0.75 mg/ml collagenase A (Roche, UK) and 1.3 mg/ml pancreatin (Sigma, UK). A combination of penicillin and streptomycin (10,000 U/ml and 10 mg/ml respectively) were added to a final concentration of 1%, and bromodeoxyuridone (100 µmol/l) was added to attenuate fibroblast proliferation. NRVMs were maintained at 37°C in a 5% CO_2_ incubator and medium changed after 48 hours (Figure 1).

**Figure 1.**
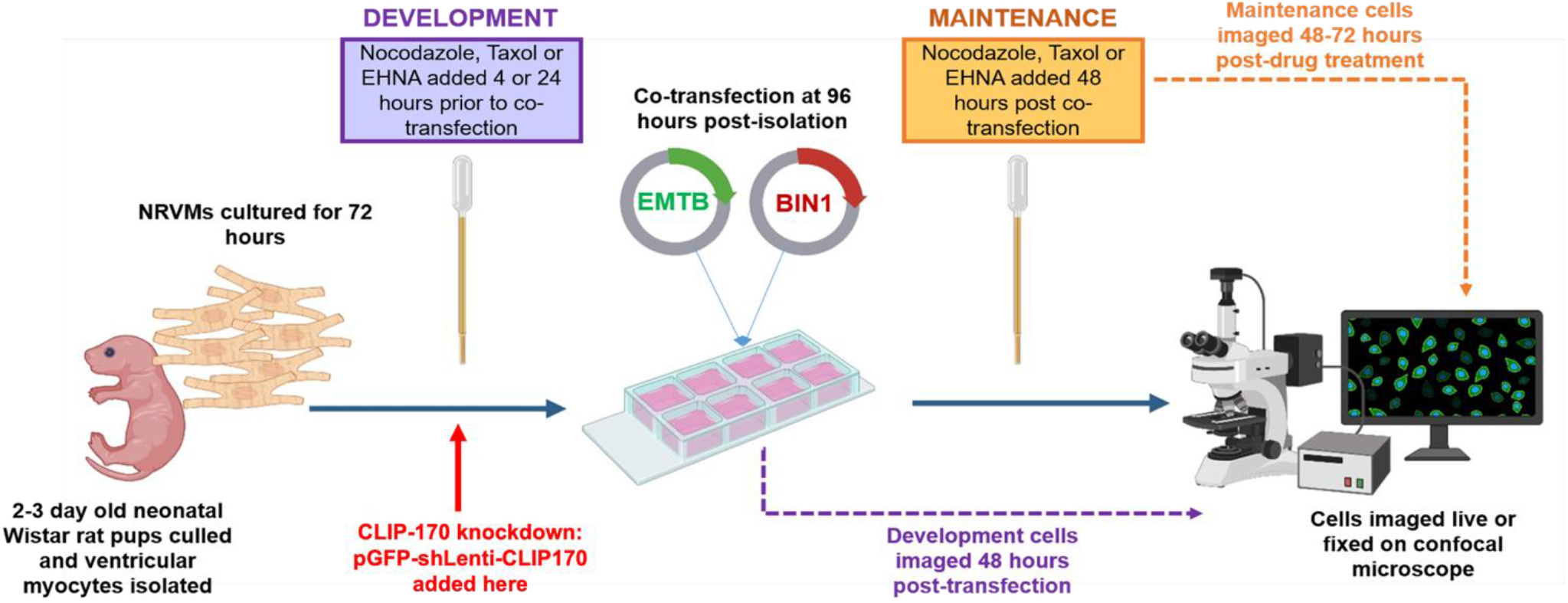
Schematic timeline of NRVM t-tubule development and maintenance studies. Purple flow chart indicates BIN1-driven tubule development experimental timeline, and orange chart indicates BIN1-driven tubule maintenance timeline. *Created with BioRender*.

### Transient transfection and transduction

To grow BIN1-driven tubules, NRVMs (1-5 days post-isolation) were transiently transfected with variant 8 of human BIN1 cloned into a pCMV6-AC-mKate2 vector (Origene Inc, USA) to induce tubule formation. An EMTB-3xGFP vector containing the microtubule binding domain of ensconsin fused to 3 x GFP molecules (Addgene, USA) was co-transfected (50:50 ratio of both plasmids) to visualise microtubules and BIN1-driven tubules within the same cell. All plasmid DNA was mixed at a 1:3 ratio with transfection reagent (Fugene 6; Promega) in reduced serum media (OptiMEM; Gibco Life Technologies, UK) and NRVMs maintained in a 5% CO_2_ incubator at 37°C for 48 hours prior to further experimentation. For CLIP-170 knockdown, NRVMs were transduced with a rat CLIP-170 shRNA construct designed against multiple splice variants, embedded in a pGFP-C-shLenti vector, and a 29-mer scrambled shRNA cassette in a pGFP-C-shLenti vector was used as a scramble control (Origene Inc, USA). For effective CLIP-170 knockdown, NRVMs were transduced with either CLIP-170 or scramble lentiviral constructs for an initial 24 hours, before BIN1-mKate transfection for a further 48 hours. A multiplicity of infection (MOI) of 5 was used for all experiments.

### Drug-induced microtubule disruption

The following drugs were used to induce microtubule disruption in NRVMs and sheep atrial myocytes: nocodazole was applied to depolymerise microtubules, Palicitaxel (taxol) was used to inhibit microtubule dynamics and erythro-9-(2-Hydroxy-3-nonyl)adenine hydrochloride (EHNA) was applied to disrupt the dynein motor protein in NRVMs only. To investigate the role of microtubules and the motor protein dynein on BIN1-driven tubule development, microtubule and dynein motor disruption was performed 48 hours prior to transient transfection and subsequent BIN1-driven tubule formation in NRVMs (Figure 1-purple flow chart). To elucidate the role of microtubules in BIN1-driven tubule and adult sheep t-tubule homeostasis, drug-induced microtubule disruption was performed 48 hours *after* transient transfection of NRVMs (Figure 1-orange flow chart), or in sheep atrial myocytes for 4 hours on the bench. For NRVMs, drugs were diluted in plating medium at the following concentrations: nocodazole, 0.8 µM; taxol, 5 µM and EHNA 10 µM. For sheep atrial myocytes, drugs were diluted in NT solution at the following concentrations: nocodazole, 20 µM and paclitaxel (taxol), 20 µM. Dimethyl sulfoxide (DMSO, 0.005%) was used as a vehicle control for all experiments. A methodological schematic of NRVM experiments is shown in Figure 1.

### T-tubule imaging and quantification

All cells in this study were imaged by confocal microscopy using a Nikon A1R+ confocal microscope, at 100 nm XY pixel dimensions and 100 nm Z-step size. Left atrial myocytes were fixed and stained with wheatgermagglutanin-488 (WGA-488) to label t-tubules and visualised using the 488 nm wavelength setting. In NRVMs, BIN1-driven tubules were visualised using the 561 nm wavelength setting (595 ± 50 nm emission) to excite the mKate fluorescent tag and at 488 nm wavelength to excite EMTB-3xGFP and visualise microtubules. Images were deconvolved, and summed stacks of 20 images (2 μM Z-projections) were thresholded using ImageJ (Fiji, National Institute of Health, USA) and backgrounds corrected. After thresholding, the following quantitative assessments of t-tubules were calculated: i) t-tubule density derived from binarised images as the fraction of pixels occupied by t-tubules relative to the planar cell area, ii) for NRVMs only, the total length of individual tubules and number of structures was determined using a ridge detection plugin. Cell size was quantified manually using the Fiji polygon tool.

### Immunofluorescence

For microtubule modulation studies, nocodazole or paclitaxel were added to sheep atrial cells for 4 hours before rapid fixation in 4% paraformaldehyde for 10 minutes. Nocodazole and paclitaxel were additionally added to the initial PBS and PFA steps, to avoid any washout effect of the drugs. Cells were subsequentially permeabilised in 0.2% Triton-X100 for 30 minutes, blocked for 1 hour in 5% normal goat serum (NGS), before a primary antibody targeting α-tubulin (1:200, ab729 Abcam) was added overnight at 4°C. A corresponding fluorescently tagged secondary antibody was then added the following day for 30 minutes at room temperature (1:200, goat anti-mouse IgG, Molecular Probes). All cells were imaged using a Nikon A1R confocal microscope as above.

### Protein Quantification

For NRVM protein extraction, cultured NRVMs were incubated in RIPA buffer containing protease and phosphatase inhibitors for 30 minutes on ice. NRVMs were then scraped and vortexed for 5 minutes, before centrifugation at 10,000 x g, 4°C and supernatants collected. Samples were prepared for separation using Novex 4 8-12% Bis-Tris gels (NuPage, Life Technologies) as described previously [58]. Following electrophoresis samples were transferred to nitrocellulose membranes (GE Healthcare, UK). Membranes were then blocked in 5% milk in TBS-T and probed with primary antibodies that detect α-tubulin (1:200, ab729 Abcam), CLIP-170 (1:50, sc-28325, Santa Cruz Biotech), JPH2 (1:100, sc-377086, Santa Cruz Biotech). HRP conjugated secondary antibodies anti-mouse IgG (1:40,000) and anti-rabbit IgG (1:40,000) were used to visualise proteins (GE Healthcare, UK). Three technical replicates were performed on each sample, and protein normalised to an internal standard control (IC) as described previously [38].

### Statistical Analysis

Distribution of all data was compared to a Gaussian distribution using the Sharipo-Wilk and Kolomogov-Smirnov normality tests (as dictated by sample size). Where data was not normally distributed, data was analysed using the appropriate non-parametric test, and data thereby presented as median values ± the interquartile (IQ) range. For normally distributed data, significance was tested with the appropriate parametric test and presented as mean values ± standard error of the mean (S.E.M). Nested statistics were performed on microtubule, BIN1-driven/ t-tubule, microtubule and western blot data from cells and repeats within the same NRVM litter or same sheep. All values were considered significant when p<0.05.

## Results

### Functional microtubules mediate BIN1-driven tubule formation

To directly investigate whether an intact microtubule network is necessary for normal t-tubule development, we first examined how microtubule depolymerisation influences BIN1-driven tubule formation. As we have previously demonstrated, NRVMs lack a mature endogenous t-tubule network yet BIN1 overexpression can initiate membrane tubulation and the formation of a nascent t-tubule network [58], which allows for the molecular dissection of pathways regulating t-tubule development and maintenance of mammalian cardiac t-tubules *in vitro* (Figure 1). NRVMs were treated and maintained in the presence of a vehicle (0.005% DMSO) or the microtubule-depolymerising agent nocodazole (0.8 µM) prior to BIN1 transfection, and BIN1-driven tubule networks analysed (Figure 2A-B).

**Figure 2.**
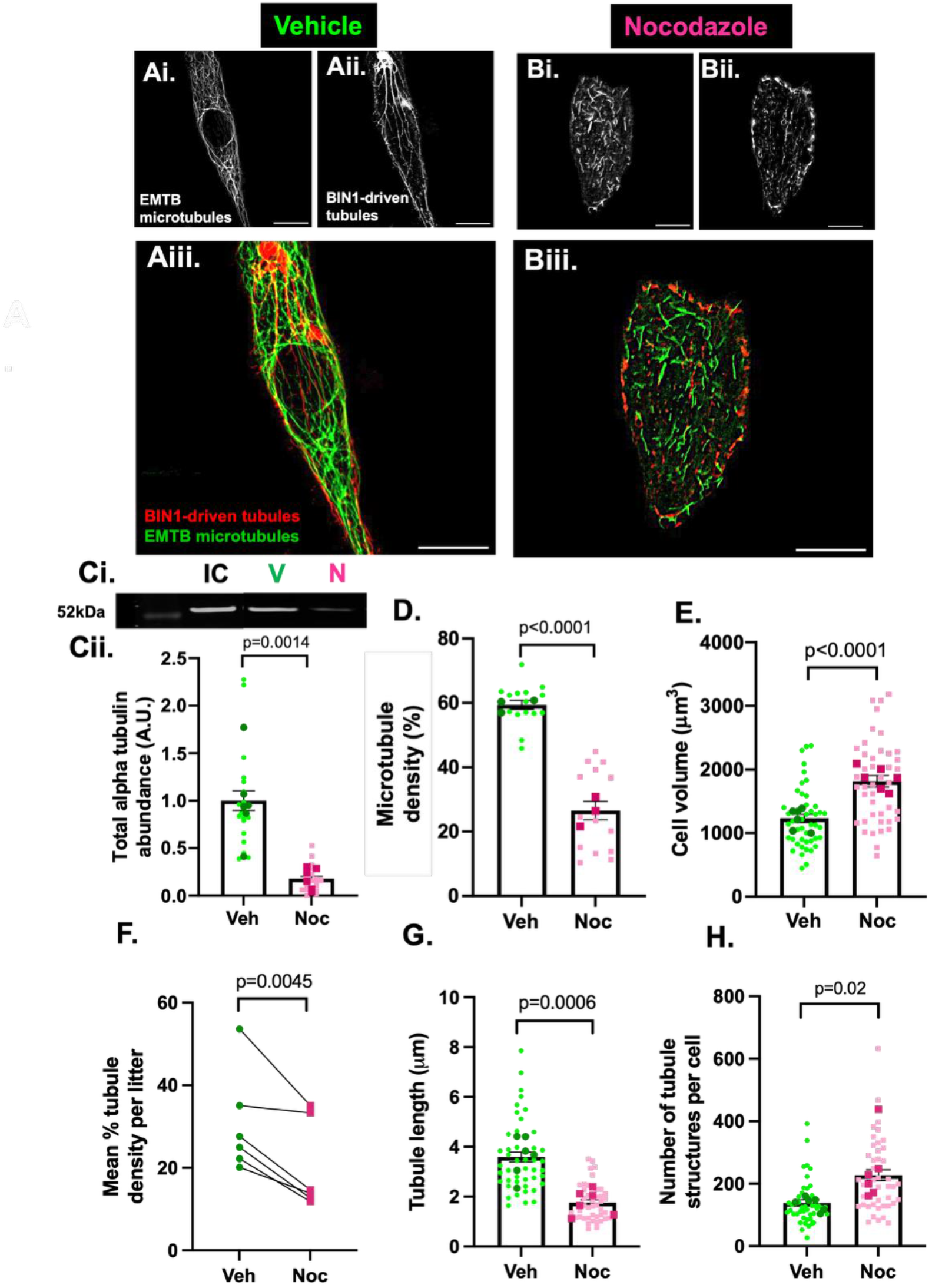
Functional microtubules mediate BIN1-driven tubule formation. Representative confocal 2 µm z-projections of vehicle (Ai-iii) and nocodazole-treated (Bi-iii) NRVMs co-transfected with BIN1-mKate (red) to form BIN1-driven tubules (Aii: vehicle-treated BIN1-driven tubule network, Bi: nocodazole-treated BIN1-driven tubule network) and EMTB-3xGFP (green) and label microtubules (Ai: vehicle-treated microtubule network, Bi: nocodazole-treated microtubule network). Merged images are shown in figure Aiii (vehicle) and Biii (nocodazole). C) Representative western blots (Ci) and mean data (Cii) of α-tubulin expression in vehicle (V) and 0.8 µM nocodazole (N) treated cells. Data normalised to total protein and internal control (IC). Cii) Total α-tubulin expression was decreased with nocodazole treatment compared to vehicle and control (p=0.0014 by nested t-test, N=6, n=3 technical repeats). D) Microtubule density is reduced 2-fold with 0.8 µM nocodazole pre-treatment (p<0.0001 by nested t-test, N<u>=3</u>, n=5-10). E) Average cell size is increased with nocodazole treatment (p<0.0001 by nested t-test, N=6, n=5-10). F) Nocodazole treatment decreases mean BIN1-driven tubule density per litter and reduces t-tubule length (G) when applied before BIN1-driven tubules are formed (p=0.0045 and p=0.0006 by unpaired and nested t-test respectively, N=6, n=5-10). H) The number of tubule structures is reduced with nocodazole treatment, indicative of t-tubule fragmentation (p=0.02 by nested t-test, N=6, n=5-10). All data plotted as mean per litter (dark points) ± SEM where light points indicate individual cell values. Scale bars 10 µm.

Protein quantification of cell lysates showed a decrease in alpha tubulin abundance following nocodazole treatment (Figure 2C; p=0.0014). Concurrently, quantitative analysis of EMTB-3xGFP-labelled microtubules demonstrated a pronounced decrease in overall microtubule density in nocodazole treated cells relative to vehicle-treated controls (Figure 2D; p<0.0001).

To account for a nocodazole-induced increase in cell size (Figure 2E; p<0.0001), BIN1-driven tubule formation was expressed as the percentage of tubules within the 2 μM Z-projection occupying the defined cell area. Mean BIN1-driven tubule density was decreased when microtubules were depolymerised with nocodazole (Figure 2F; p=0.0045). Further analysis of the BIN1-driven tubule networks in vehicle and nocodazole-treated cells revealed that tubule length was reduced with nocodazole treatment (Figure 2G; p=0.0006) and the number of structures was increased (Figure 2H; p=0.02), indicative of decreased BIN-driven tubule elongation coupled with enhanced tubule fragmentation. Interestingly, microtubule stabilisation with taxol did not alter BIN1-driven tubule development (Supplemental Figure 3). Taken together, these data delineate a profound requirement for functional microtubules in BIN1-driven tubule formation and elongation.

### Microtubules mediate BIN1-driven tubule formation via CLIP-170 and dynein-driven elongation

Given the requirement of functional microtubules in BIN1-driven tubule formation, we next sought to further explore the underlying mechanism for tubule elongation. We investigated the requirement for the motor protein dynein and the plus-end-tracking protein CLIP-170 during tubule formation.

#### Dynein facilitates tubule elongation

To determine the involvement of the molecular motor protein dynein in BIN1-driven tubule elongation, we used EHNA (10 µM) to inhibit dynein-driven retrograde movement during culture prior to and continuously during BIN1 overexpression and tubule formation (Figure 3A-B). Despite no change in cell volume (Figure 3Ci), BIN1-driven tubule density and length were decreased with no change in the overall number of detected BIN1-driven tubule structures during dynein inhibition (Figure 3Cii-iv; p=0.005-p<0.0001). Our data therefore suggests that BIN1-driven tubules fail to elongate during development when inwardly directed, dynein-driven forces are perturbed. EHNA does however have off target effects via inhibition of PDE2. To confirm a lack of involvement of PDE2 inhibition on BIN1-driven tubule formation the experiment was repeated with the PDE2 inhibitor BAY-6550. PDE2 inhibition (BAY-6550) did not alter BIN1-driven tubule density, length or number (Supplemental Figure 1) confirming the EHNA effect is not driven by PDE2 inhibition. Together, these data suggest a role for dynein-driven retrograde trafficking and/or motor protein forces, in regulating BIN1-driven tubule elongation during development.

**Figure 3.**
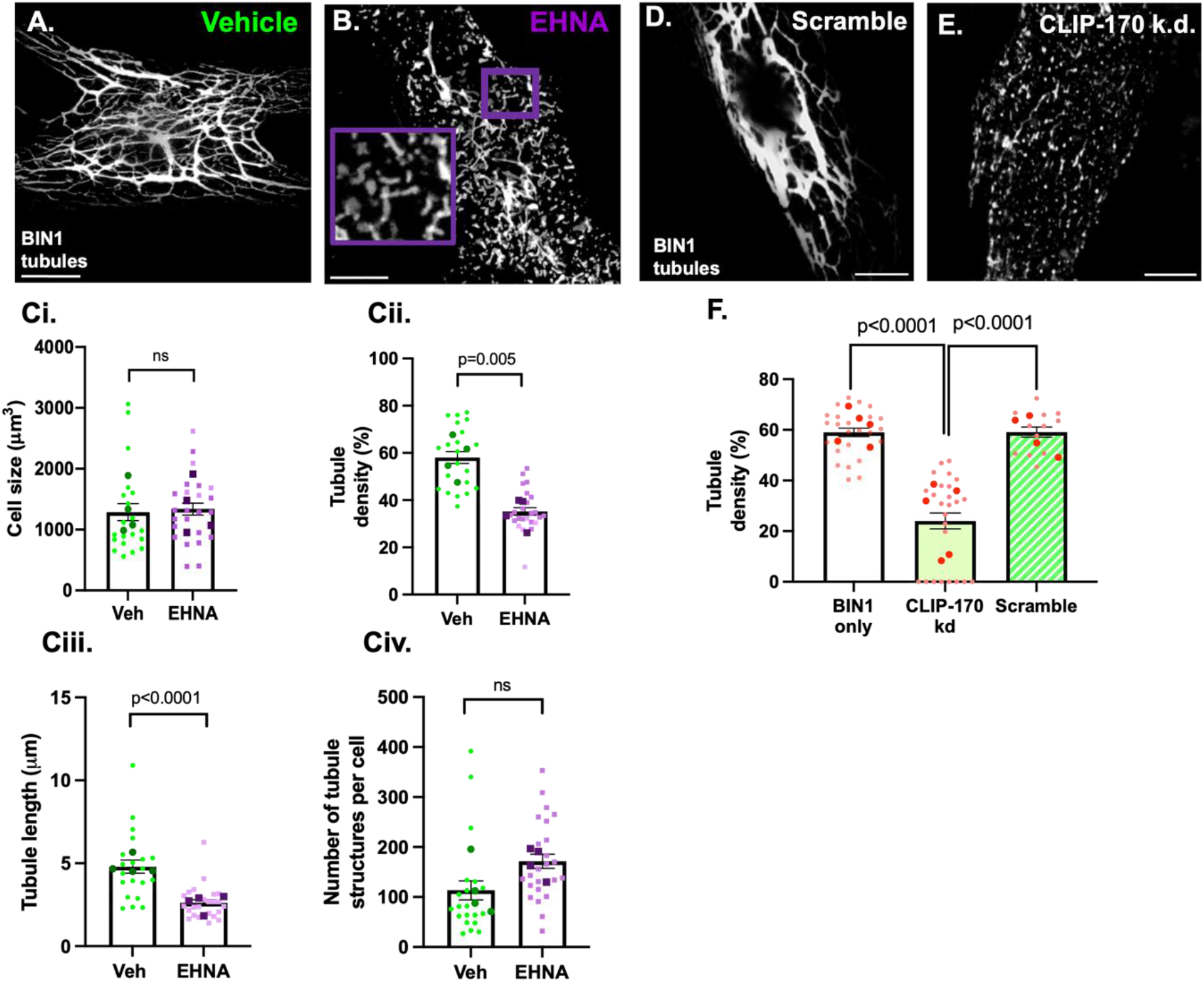
Microtubules mediate t-tubule elongation via dynein-driven movement and BIN1-CLIP-170 interactions. Representative confocal 2 µm z-projections of the BIN1-driven tubule network in vehicle (A) and 10 uM EHNA-treated (B) NRVMs. Ci) Cell size is unchanged with EHNA pre-treatment. Cii) EHNA treatment decreases BIN1-driven tubule density and reduces tubule length (Ciii) when applied before BIN1-driven tubules are formed, (p=0.005-p=<0.0001 by nested t-test, N=6, n=15-18). Civ) The number of tubule structures detected per cell is unchanged with EHNA pre-treatment. D-E) Representative confocal 2 µm z-projections of the BIN1-driven tubule network in NRVMs transduced with a pGFP-C-shLenti-CLIP170 vector to achieve CLIP-170 knockdown (D) or scramble control (E). F) CLIP-170 knockdown decreases BIN1-driven tubule density compared to scramble and BIN1 controls (both p<0.0001 by Kruskal-Wallis 2-way ANOVA, N=5, n=4-7). All data plotted as mean per litter (dark points) ± SEM where light points represent individual cell values. Scale bars 10 µm.

#### The microtubule +TIP protein CLIP-170 is required for BIN1-driven tubule development

To investigate the role of CLIP-170 in tubule biogenesis, cells were transduced with a lentiviral GFP-tagged CLIP-170 construct or scramble control, and subject to CLIP-170 knockdown or placebo prior to BIN1 transfection and BIN1-driven tubule development. Figures 3D&E show the formation of a BIN1-driven tubule network in the absence and presence of CLIP-170 KD (Figure 3D&E). The resulting effects on tubule formation were quantified, revealing a decrease in BIN1-driven tubule formation upon CLIP-170 knock down compared to the scrambled control (Figure 3D-F). Thus, we show CLIP-170 is required for normal BIN1-driven tubule formation.

### Functional and dynamic microtubules mediate BIN1-driven tubule homeostasis

#### Microtubules and dynein are required for normal BIN1-driven tubule homeostasis

Given the importance of microtubules for BIN1-driven tubule development we next sought to understand if and how microtubules influence the maintenance of new BIN1-driven tubules. NRVMs with existing BIN1-driven tubules and an EMTB-3xGFP labelled microtubule network were treated with nocodazole (0.8 µM), paclitaxel (5 µM) or EHNA (10 µM) to depolymerise microtubules, stabilise microtubules or inhibit dynein-driven movement respectively, and effects on existing BIN1-driven tubules were determined (Figure 4A-D).

**Figure 4.**
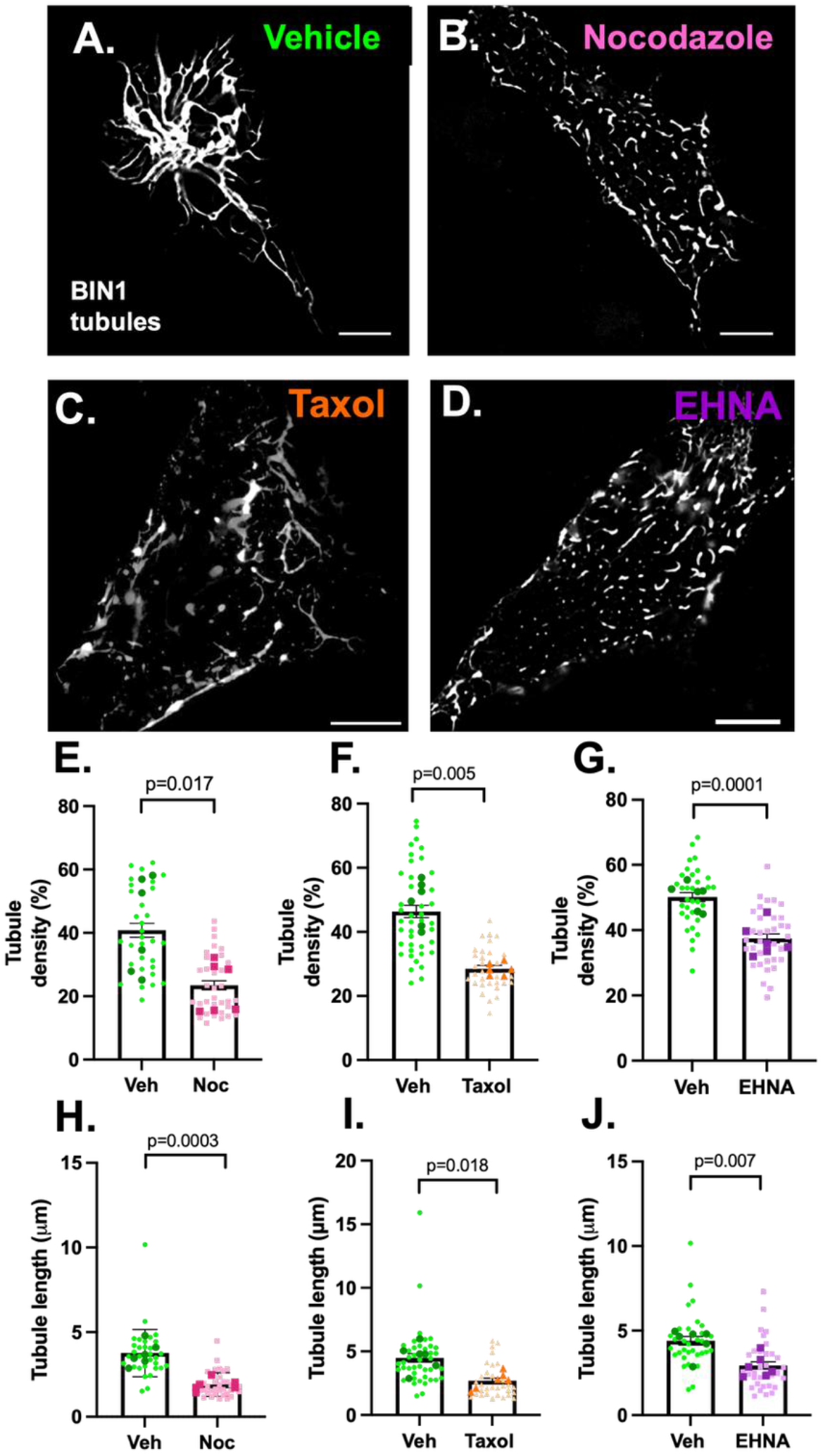
Functional and dynamic microtubules mediate BIN1-driven tubule homeostasis and elongation, alongside the molecular motor protein dynein. Representative confocal 2 µM z-projections of BIN1-driven tubules in NRVMs treated with vehicle (A), nocodazole (B), taxol (C) and EHNA (D) 48 hours post-BIN1-driven tubule formation. E-G: T-tubule density is decreased with nocodazole (E; p=0.017 by nested t-test, N=6, n=5), taxol (F; p=0.005 by nested t-test, N=6, n=5-7) and EHNA treatment to inhibit dynein (G; p=0.046 by nested t-test, N=5-7). H-J: Nocodazole treatment (H; p=0.0003 by nested t-test, N=6, n=5-7), taxol treatment (I; p=0.018 by nested t-test, N=6, n=5), and dynein inhibition with EHNA (J; p=0.007 by nested t-test, N=6, n=5) all reduced the length of tubules when applied for 48 hours post-BIN1-driven tubule formation. All data plotted as mean per litter (dark points) ± SEM where light points represent individual cell values. Scale bars 10 µm.

Interestingly, microtubule depolymerisation with nocodazole, confirmed by a reduction in microtubule density (Supplemental Figure 2A-D), markedly decreased BIN1-driven tubule density in NRVMs (Figure 4E; p=0.017) suggesting functional microtubules mediate normal BIN1-driven tubule homeostasis. Moreover, inhibiting microtubule dynamics with paclitaxel which increased the abundance of acetylated, stabilised microtubules (Supplement Figure 3H, p=0.0037), also decreased pre-existing tubule density (Figure 4F; p=0.005), signifying that dynamic microtubules also play an important role in BIN1-driven tubule homeostasis despite having no quantifiable impact on BIN1-driven tubule development in NRVMs (Supplementary figure 2A-F). Likewise, inhibition of the molecular motor dynein with EHNA perturbed pre-existing tubule turnover by decreasing density and length (Figure 4G&J; p=0.0001), indicative of reduced dynein-dependent tubule extension at the tubule membranes. All treated groups exhibited a decrease in pre-existing BIN1-driven tubule length, suggesting decreased length underlies the decrease in BIN1-driven tubule density and confirms a requirement for dynamic, functional microtubules and retrograde, dynein-driven motor protein for maintaining elongated BIN1-driven tubules (Figure 4H-J; nocodazole: p=0.0003, paclitaxel: p=0.018, EHNA: p=0.007). However, only microtubule depolymerisation with nocodazole caused pre-existing tubules to fragment (Supplemental Figure 2H; p=0.0043), and in all groups cell size remained unchanged with treatment (Supplemental Figure 2E-G). Interestingly, no change in JPH2 expression was observed with nocodazole or paclitaxel treatment, suggesting BIN1-driven tubule breakdown is microtubule – not JPH2 – mediated (Supplemental Figure 3Gi-ii).

Taken together, these data suggest a distinct requirement for functional, dynamic microtubules and a role for dynein-dependent motor forces, in BIN1-driven tubule homeostasis.

#### Functional and dynamic cardiac microtubules mediate t-tubule homeostasis in adult sheep atrial myocytes

Finally, to confirm whether atrial t-tubule maintenance in adult cardiac myocytes also requires microtubules, we investigated the effects of microtubule depolymerisation and stabilisation on sheep atrial t-tubules. Freshly isolated left atrial myocytes from control sheep were incubated with a vehicle control (0.005% DMSO), nocodazole or taxol (both 20 µM) for 4 hours at room temperature, before fixation and subsequent immunolabelling to visualise the microtubule and t-tubule networks (Figure 5A-F). Figure 5A and 5B show a typical atrial myocyte microtubule and t-tubule network respectively after 4 hours. Similar data is shown following 4 hours incubation with either nocodazole (Figure 5C&D) or taxol (Figure 5E&F).

**Figure 5.**
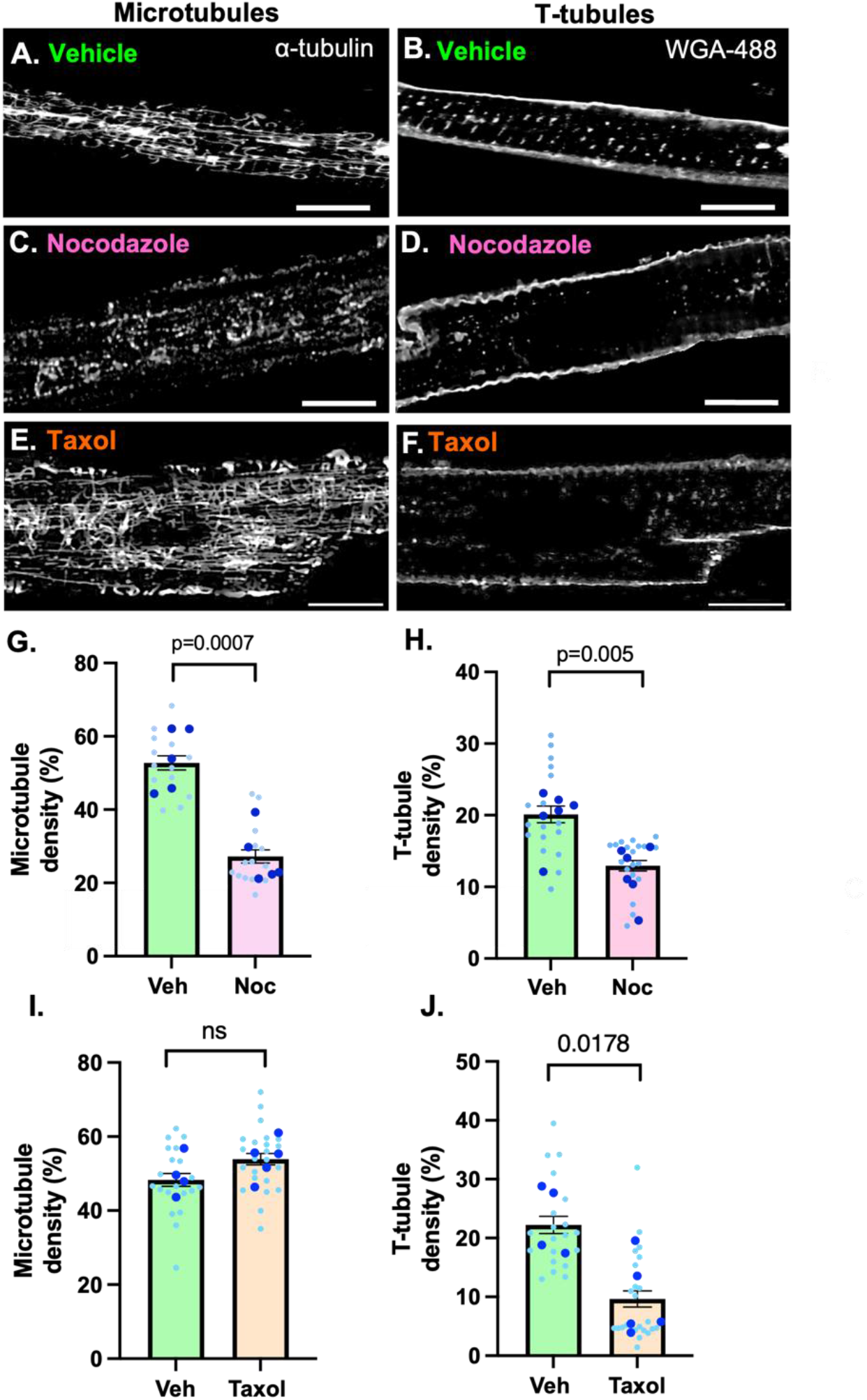
Functional and dynamic microtubules modulate t-tubule homeostasis in the normal atria. Representative confocal 2 µM z-projections of the microtubule network (A,C,E) labelled with α-tubulin) and t-tubule network (B,D,F; stained with WGA-488) in control sheep atrial myocytes treated with vehicle (A and B), 20 µM nocodazole (C and D) and 20 µM taxol (E and F). G-H) Microtubule density is decreased with nocodazole treatment (p=0.03 by nested t-test, N=5, n=1-6 cells per animal) and t-tubule density is also reduced (p=0.005 by nested t-test, N=6, n=17-19). I-J) Microtubule density is unchanged with taxol treatment (I) but t-tubule density is reduced (p=0.0178 by nested t-test, N=4-5, n=2-7 cells per animal). Data plotted as mean per animal (dark points) ± SEM where light points represent individual cell values. Scale bars 10 µm.

**Figure 6.**
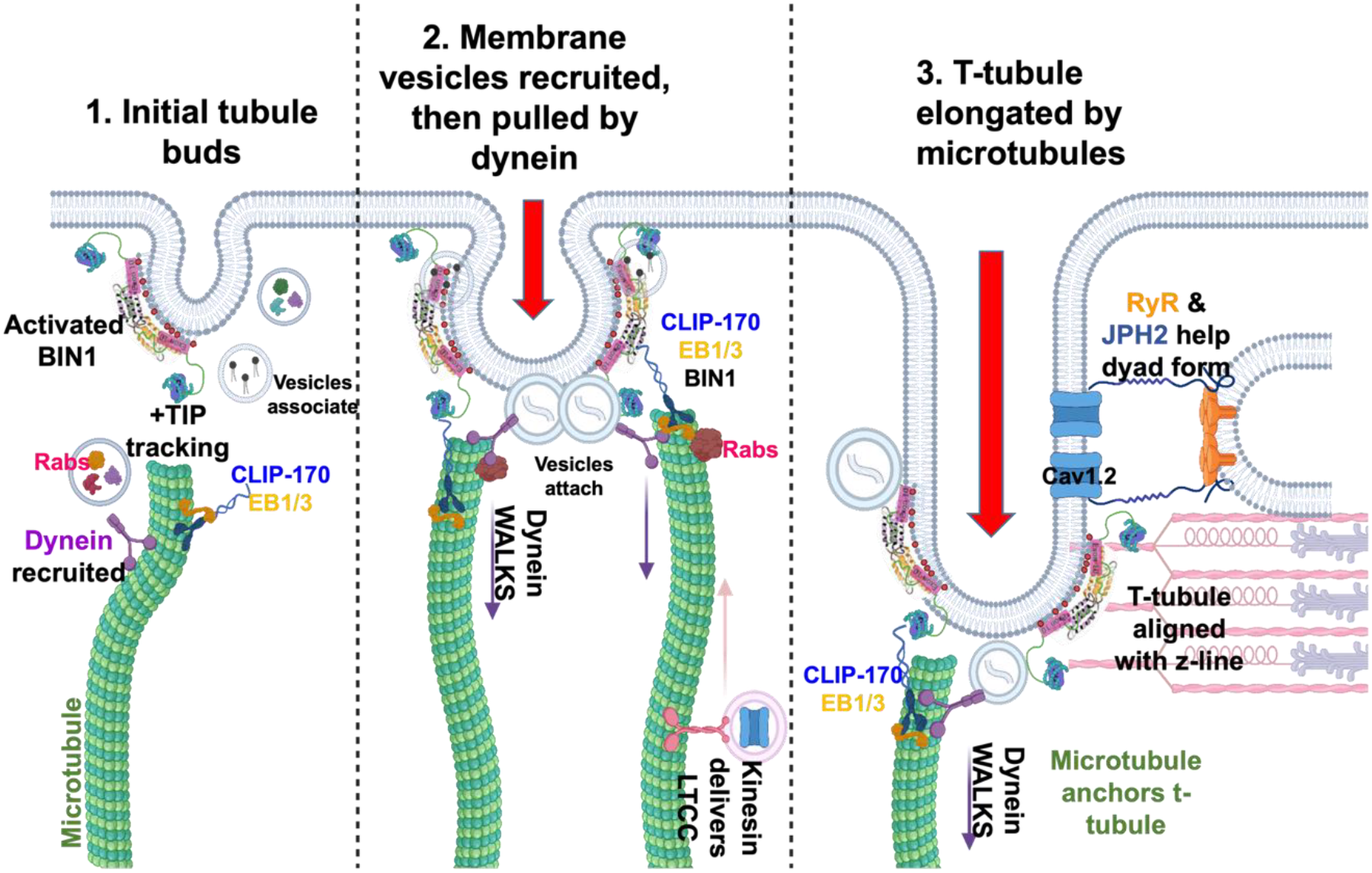
Schematic diagram depicting microtubules in t-tubule elongation. Microtubules associate with the budding tubule via TIP+ tracking and dynein is recruited with the help of Rab proteins. BIN1 is activated by dynamin and is associated with CLIP-170 and EB1/3 to anchor microtubule, and dynein attaches to endocytic membrane vesicles. Dynein walks down the microtubule, pulling the budding t-tubule with it, and kinesin walks towards the +TIP to help deliver LTCCs. Microtubules guide the t-tubule and anchor to the z-line, where RyR and JPH2 along with BIN1, help form the dyad and a stable, elongated tubule. *Created in Biorender*.

Quantitative analysis of both microtubule and t-tubule networks revealed that nocodazole-induced microtubule depolymerisation versus vehicle control (Figure 5G; p=0.0007) which resulted in a concurrent and profound loss of atrial t-tubules in control myocytes (Figure 5H; p=0.005), confirming sheep atrial t-tubule integrity is dependent on functional microtubules in adult cardiac myocytes. Furthermore, stabilising microtubules with paclitaxel did not alter microtubule density (Figure 5I) but decreased atrial t-tubule density (Figure 5J; p=0.0178). Microtubule dynamics therefore play an important role in t-tubule turnover in adult atrial myocytes.

Collectively, this study provides novel mechanisms for microtubule-mediated tubule development and tubule turnover processes, demonstrating a new and exciting functional link between microtubule and t-tubule networks in health.

## Discussion

This study identifies the cardiac microtubule network as an active determinant of t-tubule formation and maintenance. Using BIN1-driven tubules in NRVMs and native t-tubules in adult sheep atrial myocytes, we show that i) t-tubule growth requires not only BIN1-dependent membrane remodelling, but also an intact microtubule network, and ii) CLIP-170-dependent plus-end anchoring and dynein-associated motor activity is also required for BIN1-driven tubule formation in NRVMs. We show iii) functional and dynamic microtubules mediate BIN1-driven tubule turnover in NRVMs alongside dynein, and iv) depolymerisation or stabilisation of the adult atrial cardiac microtubule network destabilised native atrial t-tubules. These data move the field beyond the concept of microtubules as passive tracks for channel delivery and support a model in which intact microtubules regulate BIN1-driven tubule growth via CLIP-170 and dynein-driven forces and support a role for functional and dynamic microtubules in t-tubule turnover.

### Microtubules mediate BIN1-driven tubule formation via CLIP-170 and dynein

A central finding is that microtubules are required during the earliest stages of BIN1-driven tubule biogenesis. NRVMs lack a mature endogenous t-tubule network yet have responsive and dynamic microtubule network, making them a useful reductionist model in which BIN1 overexpression via transient transfection can initiate membrane tubulation [59, 60]. In this setting, low dose nocodazole markedly reduced tubule density and length, while increasing the number of detected structures. This phenotype is most consistent with failed elongation and fragmentation of nascent tubules, rather than a simple reduction in BIN1 expression or cell size and indicates that BIN1-mediated membrane bending alone is insufficient to generate an extended tubule network without a functional microtubule cytoskeleton.

These findings extend current models of t-tubule biogenesis. Cardiac t-tubules are proposed to arise from BIN1-enriched membrane domains that recruit lipids, curvature-generating proteins and vesicular cargo to form early dyadic structures [39–41, 61]. Previous work has shown that BIN1 can fold cardiac membrane, organise LTCC-containing microdomains and tether dynamic microtubules to deliver Cav1.2 to t-tubule membranes [22, 23]. In non-muscle cells, BIN1-induced membrane tubulation also depends on the microtubule plus-end protein CLIP-170, supporting the idea that BIN1 can act as a membrane anchor for growing microtubule ends [24, 56]. Our CLIP-170 knockdown experiments place this interaction in a cardiac tubule-forming context: loss of CLIP-170 reduced BIN1-driven tubule formation, indicating that microtubule plus-end capture is a functional step in cardiac tubule initiation. We therefore suggest that BIN1 acts not only as a membrane-bending scaffold but may also act as a cortical docking site that couples nascent tubule membranes to dynamic microtubule plus ends, enabling directed delivery of membrane and dyadic cargo during t-tubule growth.

Our data further suggest that microtubule-dependent tubule formation likely requires a microtubule motor protein component. Inhibition of dynein reduced BIN1-driven tubule density and length without increasing the number of structures, consistent with impaired inward extension of nascent tubules rather than enhanced fragmentation. This aligns with evidence from endocytic systems in which microtubule motors power membrane curvature and elongate developing endocytic tubules [55]. This is consistent with work in isolated adult rat ventricular myocytes, within which acute EHNA-treatment (30 μM for 120 minutes) perturbed dynein-driven SR motility [62]. Therefore, in cardiomyocytes, dynein may contribute to tubule development by transporting tubule-associated cargo, by organising the position and tension of microtubule ends at BIN1-positive membranes, or by directly supporting inward membrane extension along the microtubule network. Although EHNA-based dynein inhibition has limitations and does not prove direct motor engagement at the tubule tip, the absence of comparable effects with selective PDE2 inhibition strengthens the interpretation that reduced dynein-dependent motility contributes to the phenotype (Supplemental Figure 1). Future live-cell imaging of BIN1, CLIP-170, dynein and microtubule plus ends will be important to distinguish whether dynein primarily drives tubule elongation, cargo delivery, or both.

### Microtubules critically regulate t-tubule turnover

We show that microtubules are also required after tubules have formed. T-tubules are increasingly recognised as dynamic membrane structures that remodel during development, disease and recovery [29, 38, 48, 54, 63]. Consistent with this concept, microtubule disruption or stabilization depleted the existing BIN1-driven tubule network in NRVMs, while dynein inhibition produced a similar loss of tubule density and length, indicating that BIN1-driven tubule maintenance requires microtubule dynamics rather than simply microtubule mass. This distinction is important because it suggests that ongoing growth, shrinkage and motor-dependent remodelling of the microtubule network are needed to maintain membrane continuity, cargo exchange and structural support at BIN1-positive tubules.

Our adult sheep atrial data provide important physiological support for this model. Within only four hours, both nocodazole and taxol reduced native atrial t-tubule density, demonstrating that even established atrial t-tubules require a functional and dynamic microtubule cytoskeleton for stability. This is consistent with recent experimental work showing that recovery of t-tubules after osmotic shock-induced de-tubulation is slowed by either microtubule disruption or stabilisation, and with physiological rodent studies linking microtubule disruption to defective trafficking of t-tubule-associated proteins such as JPH2 [52, 53]. However, in our experiments in NRVMs, JPH2 abundance was not altered by nocodazole or paclitaxel (Supplemental Figure 3G), suggesting that the early loss of tubule structure observed here is unlikely to be explained simply by reduced total JPH2 expression. Moreover, recent work has demonstrated that cardiac t-tubules are actively removed through PKC–PKD–NFκB-dependent macropinocytosis during the disease state, further establishing t-tubule loss as a regulated membrane turnover process rather than passive structural degeneration [64]. However, the upstream cytoskeletal mechanisms that spatially regulate this membrane retrieval and its subsequent intracellular trafficking remain unknown in this study. Instead, our findings here point toward a more direct requirement for microtubule dynamics, BIN1-associated membrane trafficking and motor-dependent membrane remodelling in maintaining the atrial t-tubule network. This may be particularly relevant in large mammalian atria, where t-tubules are well developed and make a major contribution to synchronised Ca^2+^ release, but are profoundly depleted in HFrEF and where JPH2 may play a different role [44, 58].

Collectively, these data support a model in which cardiac t-tubules are sustained by a dynamic BIN1-microtubule axis. In this model, BIN1-rich membrane domains recruit CLIP-170-positive microtubule plus ends to initiate tubule formation, while dynein forces and ongoing microtubule turnover support inward elongation and maintenance of the tubule membrane. This framework provides a mechanistic link between two hallmarks of cardiac remodelling: t-tubule loss and microtubule dysfunction. While future studies should define the live dynamics of BIN1, CLIP-170, dynein and t-tubule membranes and determine the functional consequences for atrial Ca^2+^ release and contraction, the present findings identify microtubules as active regulators of t-tubule architecture and raise the possibility that restoring appropriate microtubule dynamics could help preserve or recover dyadic structure in cardiac disease.

## Supplemental Materials

**Supplemental figure 1.**
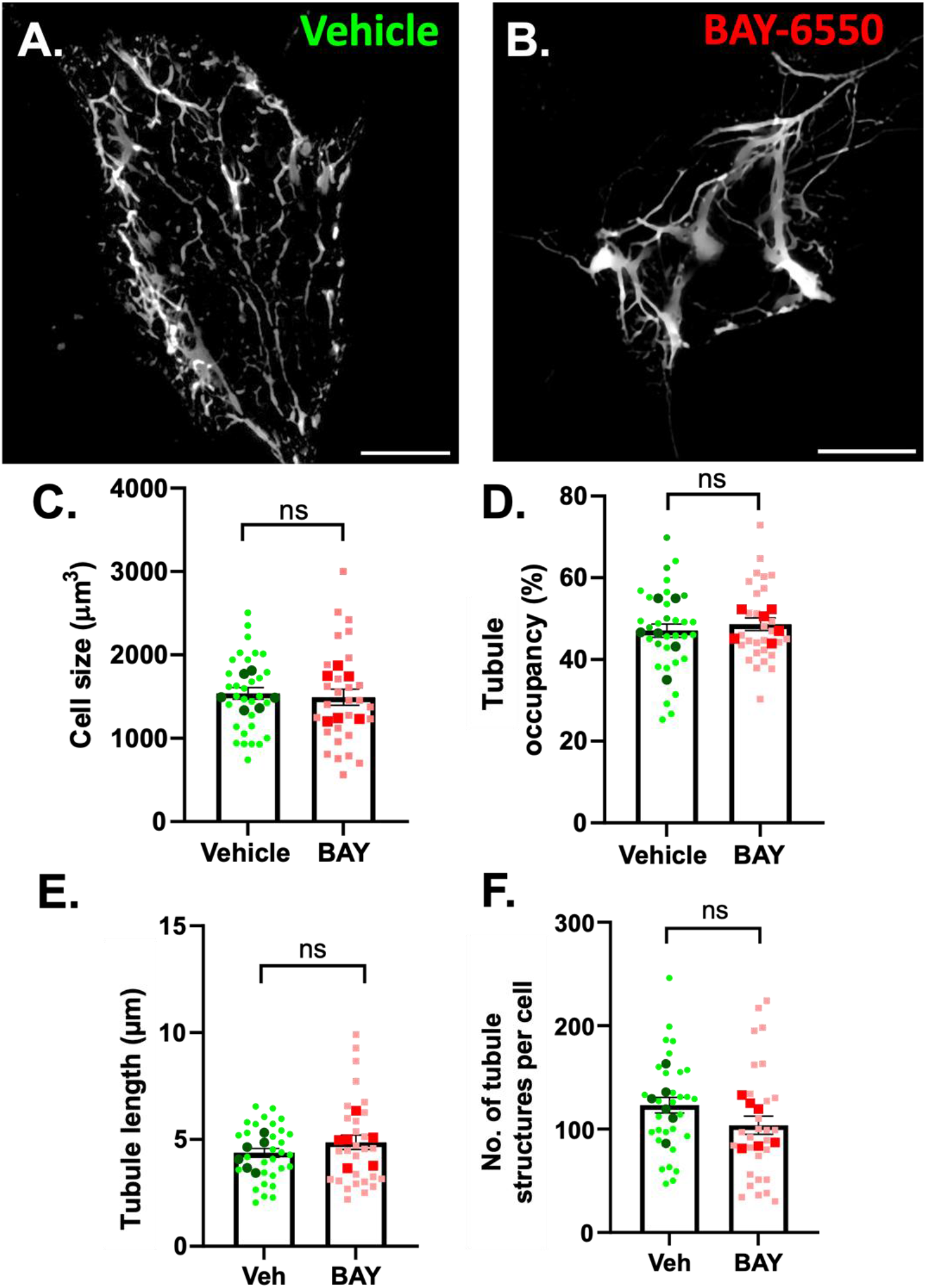
PDE2 inhibition with BAY-6550 does not alter BIN1-driven tubule development. Representative 2 μM Z-projections of NRVMs treated with vehicle (A) or 10 μM BAY-6550 to inhibit PDE2, 48 hours prior to BIN1-driven tubule formation. C-F) Pre-treatment with BAY-6550 did not perturb BIN1-driven tubule density, length or number, nor did it alter cell size (N=6 litters, n=3-5 cells per litter). All data plotted as mean per litter (dark points) ± SEM where light points indicate individual cell values. All scale bars 10 μM.

**Supplemental figure 2.**
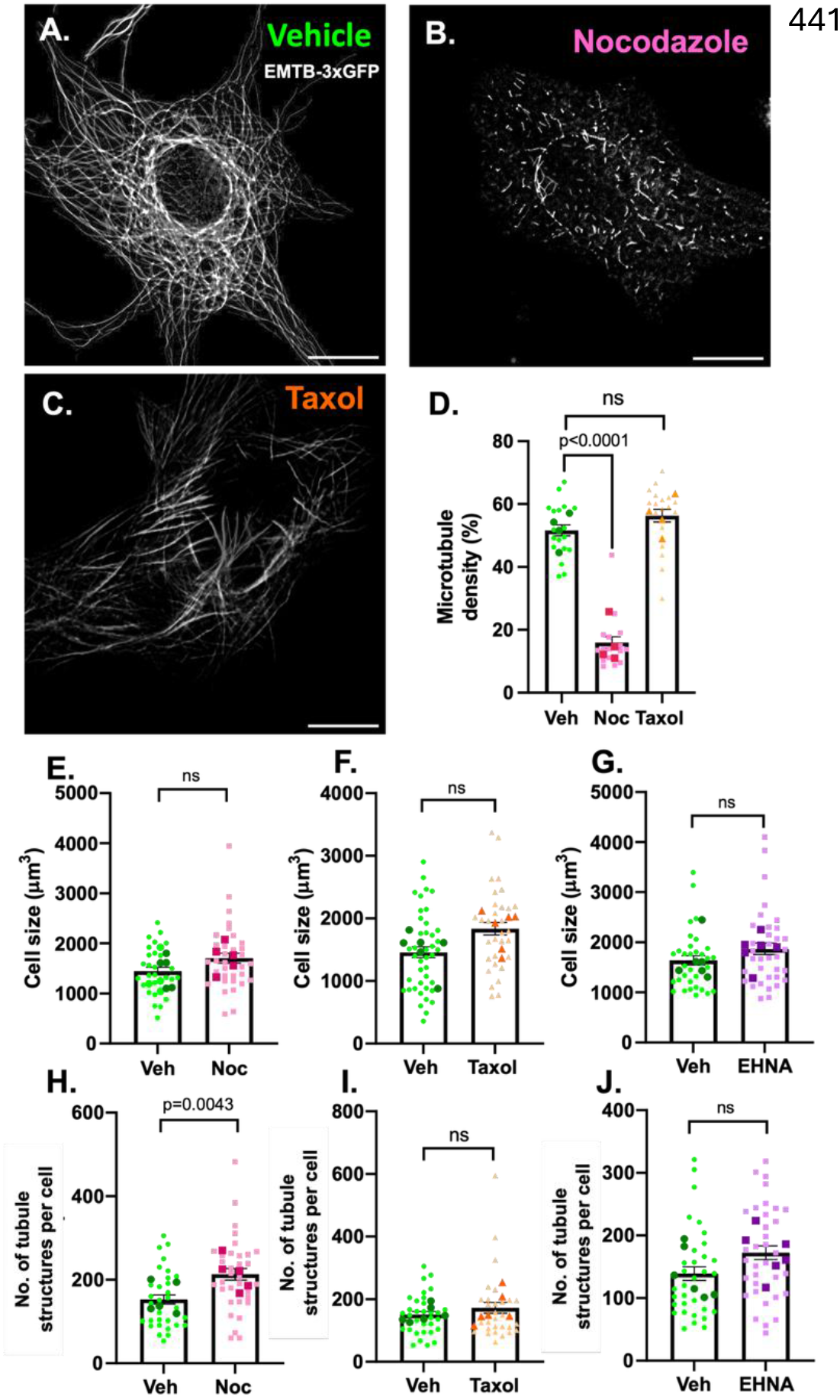
Microtubule density, cell size and number of structures for BIN1-driven maintenance studies in NRVMs. A-C) Representative 2 μm z-projections of microtubule networks labelled with EMTB-3xGFP and treated with vehicle (A; 0.005% DMSO, 0.8 μM nocodazole (B), or 5 μM taxol. D) Microtubule density is reduced with nocodazole treatment compared to vehicle control (p<0.0001 by nested t-test, N=4, n=4-5), but unchanged with taxol treatment. E-G) Cell size measurements were unchanged with nocodazole (E), paclitaxel (F) or EHNA (G) treatment (N=5-6 litters, n=5-7 cells per litter). H-J) The number of t-tubule structures was increased with nocodazole treatment (H; p=0.0043 by nested t-test, N=6, n=5), but no change in structures was observed with taxol or EHNA. All data plotted as mean per litter (dark points) ± SEM where light points represent individual cell values.

**Supplemental figure 3.**
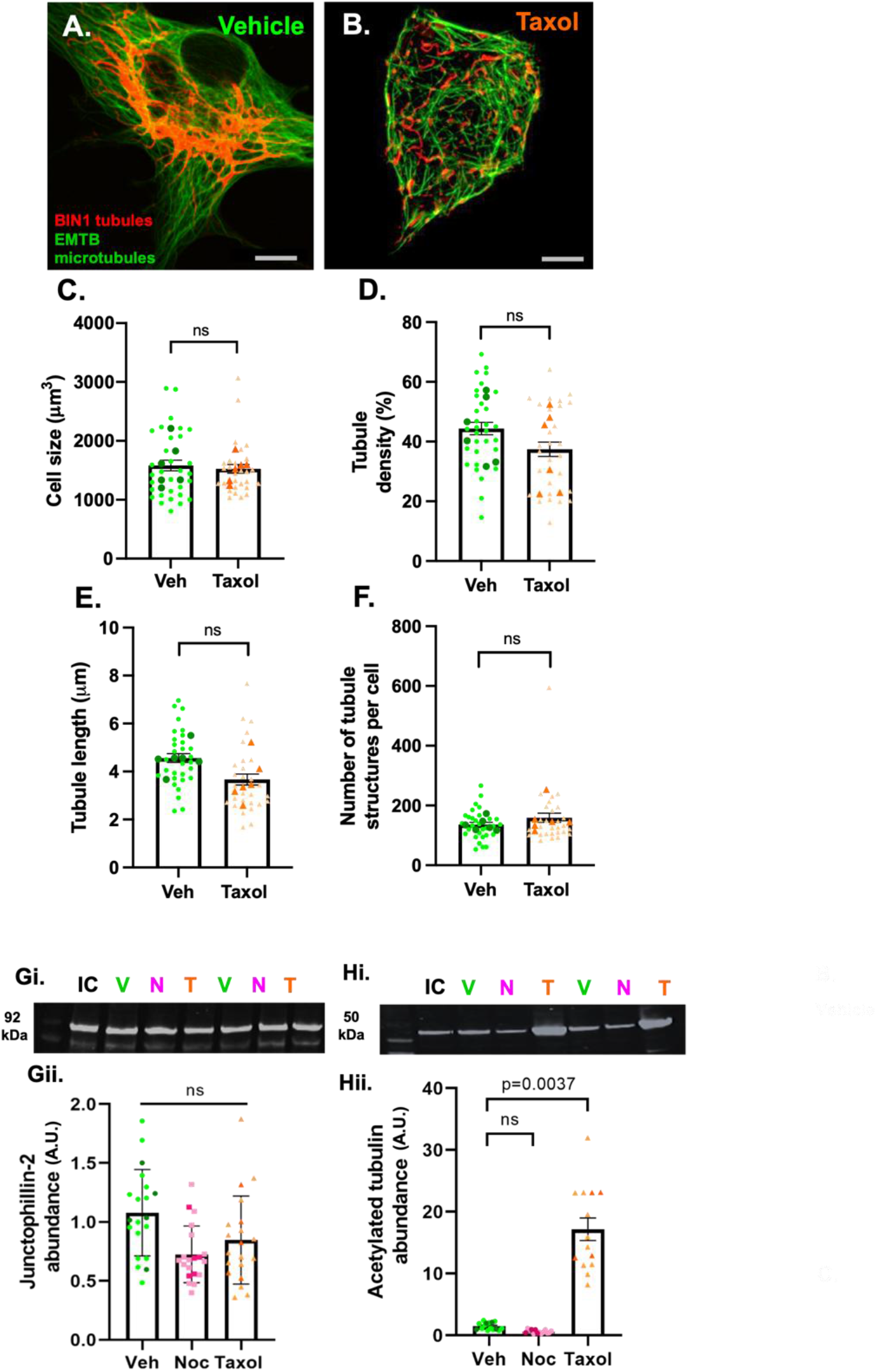
Effects of microtubule stabilisation on BIN1-driven tubule growth. Representative confocal 2 µm z-projections of vehicle (A) and 5 μM taxol-treated (B) NRVMs co-transfected with BIN1-mKate (red) and EMTB-3xGFP (green) to drive tubule formation and label microtubules respectively. C-F) No alterations to cell size (C), or BIN1-driven tubule occupancy (D), length (E) and structure (F) were observed with taxol pre-treatment. Gi-H i) Representative western blots (G-Hi) and mean data (G-Hii) of Junctophillin-2 (G) and acetylated tubulin (H) abundance in vehicle (V), 0.8 µM nocodazole (N) and taxol (T) treated cells. Data normalised to total protein and internal control (IC). No change in JPH2 abundance was observed with nocodazole or taxol treatment (N=5 litters, n=3 technical repeats). H) Acetylated tubulin abundance was markedly increased with taxol treatment (p=0.0037 by nested t-test, N=4 litters, n=3 technical repeats). All data plotted as mean per litter (dark points) ± SEM where light points represent individual cell values. Scale bars 10 µm.

## Notes

### Competing Interest Statement

The authors have declared no competing interest.

### Funding

- British Heart Foundation: FS/19/60/34899A, PG/23/11642

